# Synchronous Caregiving from Birth to Adulthood Tunes Humans’ Social Brain

**DOI:** 10.1101/2020.03.09.974659

**Authors:** Adi Ulmer-Yaniv, Roy Salomon, Shani Waidergoren, Ortal Shimon-Raz, Amir Djalovski, Ruth Feldman

## Abstract

Mammalian young are born with immature brains and rely on the mother’s body and caregiving behavior for maturation of neurobiological systems that sustain adult sociality. However, the parent-child precursors of humans’ social brain are unknown. We followed human neonates, who received or were deprived of maternal bodily contact, to adulthood, repeatedly measuring mother-child interactive synchrony. We tested the neural basis of empathic accuracy in adulthood and utilized multivariate techniques to distinguish brain regions sensitive to others’ distinct emotions from those globally activated by the vicarious stance. A network comprising the amygdala, insula, and temporal pole underpinned empathic accuracy, which was shaped by mother-child synchrony across development. Synchronous experiences with mother or father in infancy impacted adults’ neural empathy, highlighting the benefits of humans’ bi-parental rearing. Findings demonstrate the centrality of synchronous caregiving across development for tuning humans’ social brain.

## Introdunction

Being born a mammal implies that the brain is immature at birth and develops in the context of the mother’s body and caregiving behavior. Infants rely on the provisions embedded in the mother’s body, such as smell, touch, heat, or movements, and the expression of caregiving behavior for maturation of neurobiological systems that sustain participation in the social world. For mammalian young, therefore, no factor is as critical to brain maturation as the dependence of the infant on its mother, and no feature of ontogeny is as consequential as proximity to mother’s body and the experience of synchronous caregiving ^1^. Extant research in animal models has shown that breeches in the mother’s continuous presence and variability in caregiving behavior carry long-term effects on brain structure and function, particularly on systems that underpin sociality, and the effects are maintained throughout life, altering the adult animal’s capacity to coordinate social bonds, manage hardships, and parent the next generation ^2,3^. However, while the human brain is slowest to mature and requires the most extended period of dependence, the long-term consequences of parenting on the human social brain are unknown. To date, no study has followed infants from birth to adulthood to test whether variability in maternal contact and synchronous caregiving longitudinally impact social brain functioning in human adults.

The human social brain integrates activity of subcortical, paralimbic, and cortical structures to sustain the multi-dimensional task of managing human social life, which requires rapid processing of social inputs, top-down interpretation of others’ intent, and coordination of the two into social action in the present moment ^4^. The social brain has undergone massive expansion across primate evolution to support humans’ exquisite social skills, communicative competencies, and mindreading capacities and it is suggested that *Homo sapiens’* success over other hominin owes to their empathic abilities to quickly identify and mentally share others’ emotional states ^5^. Such multifaceted empathy, which integrates automatic identification of others’ emotions with the vicarious stance, marks a fundamental achievement of the social brain that enabled humans to coordinate actions for survival, jointly execute collaborative goals, and fine-tune communicative signal systems. The integrated neural system that sustains empathy combines nodes implicated in rapid assessment of salience cues with regions underpinning top-down affect sharing ^6^. Yet, while the brain basis of empathy defines a core environment-depended network tuned in mammals by parental care, the relational precursors of the neural empathic response have not been studied in humans.

Interaction synchrony, the coordinated exchange between parents and infants where parent matches the infant’s nonverbal signals online, is a prototypical experience that prepares infants to life with others and sharpens social-emotional competencies. Through moment-by-moment responsivity to infant communications and imitation, parents orient children to social moments, practice rapid assessment of distinct emotional states, and, over time, enable children to give meaning to others’ social actions, fine-tuning the social brain and its capacity to sustain empathy ^7^. Mother-child synchrony undergoes significant maturation across child development and evolves from non-verbal matching to verbal dialogue that acknowledges others’ emotions, engages multiple perspectives, and reflects on feelings; still, the synchronous dialogue retains its basic rhythms from infancy. Mother-infant synchrony is individually stable across childhood and shapes empathic abilities in adolescence ^8^.

The development of synchrony is highly sensitive to initial conditions. Conditions that compromise maternal-infant bonding bear long-term negative consequences for the synchronous exchange and, consequently, for maturation of human social abilities ^9,10^. When infants are born prematurely and are deprived of maternal bodily contact, the development of synchrony is halted and socioemotional competencies compromised. Notably, when we provided structured maternal-infant skin-to-skin contact (Kangaroo Care, KC) to premature neonates during the postpartum period of separation, the intervention improved not only interaction synchrony but also the functioning of regulatory support systems, such as circadian rhythmicity, autonomic maturity, stress responsivity, and exploratory behavior, the same systems impacted in young mammals by contact with the maternal body ^11^.

What may be the effects of maternal-newborn bodily contact and synchronous caregiving experienced across development on adults’ social brain? Utilizing our unique cohort, we imaged the neural empathic response in adults reared under different initial conditions, including infants born at full-term (FT), preterm infants receiving kangaroo contact (KC), and demographically- and medically-matched preterm infants receiving standard incubator care (SC) who were followed in our lab for two decades (Fig. 1A). We focused on the neural basis of empathic accuracy, the capacity to detect and affectively share others’ distinct emotions, considered a core feature of empathy ^12^. Employing a validated fMRI paradigm that exposed participants to others’ distinct emotions (joy, sadness, distress) and activated the vicarious position ^13^ (Fig. 1B), we used Representational Similarity Analysis (RSA), a multivariate brain pattern analytic technique, to discover the brain systems that uniquely respond to different emotions from those globally activated to affect sharing. RSA compares the similarity of neural patterns across different conditions to characterize the representational geometry of empathy to different emotions ^14^. Critically, we examined how mother-child synchrony, repeatedly observed from infancy to adulthood, tunes the accuracy of the neural empathic response in human adults.

**Fig. 1.**
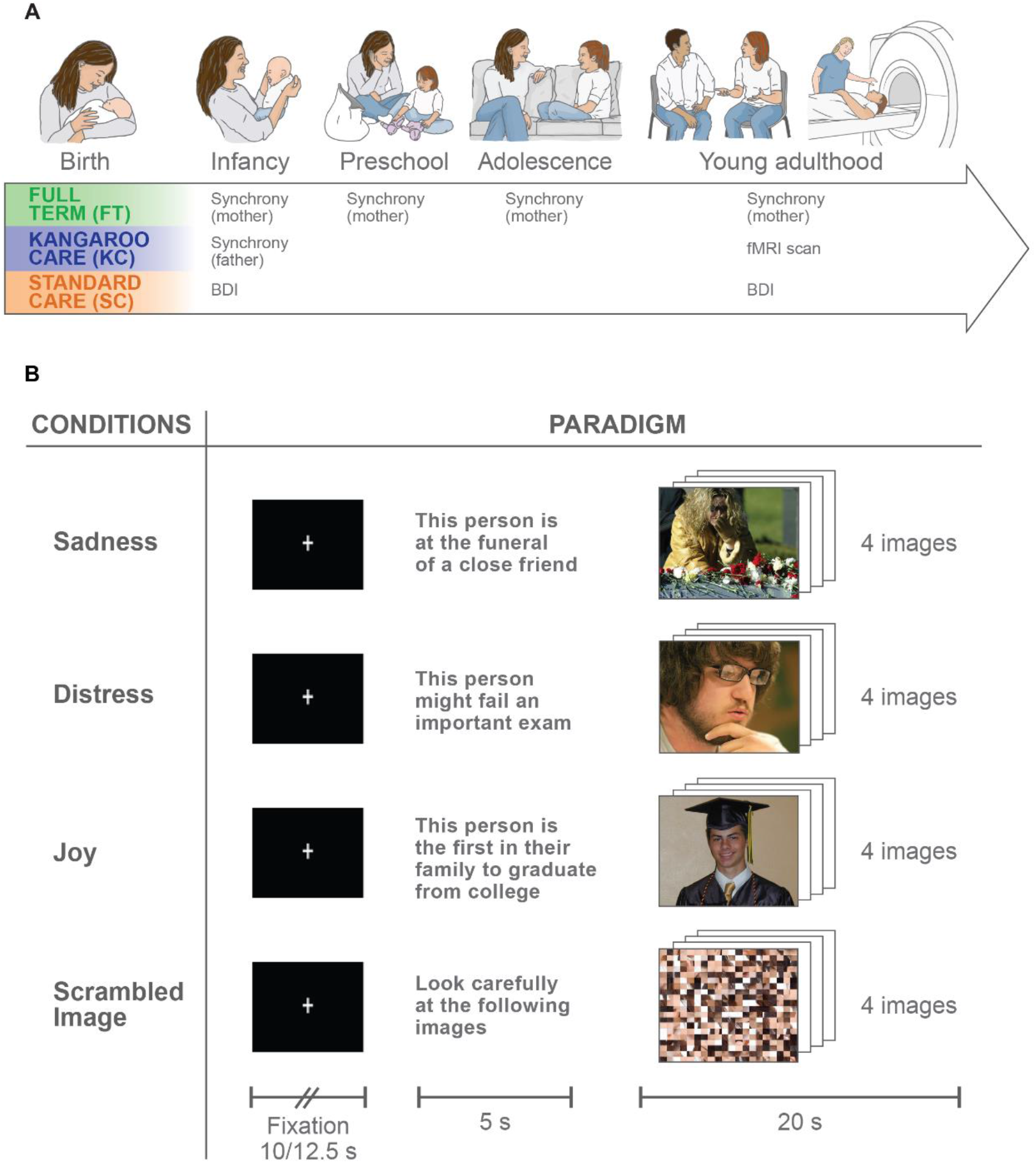
Birth-to-adulthood longitudinal study design and fMRI paradigm. (**A**) Three cohorts of infants and parents recruited at birth: Full-term infants (FT) and two matched neurologically-intact premature infants randomly assigned to Kangaroo Care (KC), infants receiving skin-to-skin contact with mother and matched controls receiving standard incubation care (SC). Mother-child interactive synchrony was assessed at 4 months (SD =1.14), 3 years (SD = 1.38), 12 years (SD = 1.62), and 20 years (SD = 2.01), and father-infant synchrony at 4 months. **(B)** fMRI “empathic accuracy” paradigm. Example illustrating a pseudorandomized-design where participants presented an emotional probe followed by 4 photos depicting it. Participants asked to empathize with protagonists and 5 blocks per condition were presented.

Using a large sample (N=97), we first examined the neural basis of empathic accuracy. Based on prior imaging studies we expected that a network including limbic regions, such as the amygdala and parahippocampal gyrus; the anterior insula, superior temporal sulcus (STS), and temporal pole (TP), found to activate empathy studies ^15,16^; and the ventromedial prefrontal cortex (VMPFC), posterior cingulate cortex, and temporal parietal junction, which comprise the default mode network (DMN), would activate when humans empathize with others’ emotions.

We then investigated which regions show differential representational geometries for empathy to distinct emotions versus regions indifferent to specific emotional content. We expected that the amygdala, which rapidly responds to variations in emotional salience, would exhibit differential activation patterns to each emotion. In parallel, we examined whether structures of the DMN, including precuneus and VMPFC, whose role in empathy has been established, would respond globally to affect sharing. Our key hypothesis was that the experience of parent-child synchrony across development, by which children practice the identification and sharing of others’ emotions, would enhance the neural differentiation of empathy to others’ distinct emotions.

## Results

To chart the long-term effects of maternal-newborn bodily contact on the development of synchrony, we first examined the maturation of *synchrony* from infancy to adulthood in the three groups. Repeated measures ANOVA revealed that *synchrony* increased considerably from infancy to adulthood across all participants (Fig. 2), highlighting its maturation from non-verbal exchange to verbal dialogue of mutuality and sharing (*F_(3,228)_*=161.95, p<0.001, η^2^_p_ =0.68). Significant group differences emerged (*F_(2,76)_*=18.58, *p*<0.001, *η^2^_p_* =0.32); FT mother-child dyads exhibited the highest *synchrony* (Mean (M)=3.58, Standard Deviation (SD)=0.70, p=0.031, p<0.001, respectively) and KC mother-child dyads showed more *synchrony* than SC dyads who received no maternal-newborn contact (KC:M=3.26, SD=0.89, p=0.007, SC: M=2.88, SD=0.88, Fig. 2). A significant interaction of group and time was revealed (*F_(6,228)_*=2.75, p=0.013, η^2^_p_ =0.068). During infancy, FT children engaged in more *synchrony* than KC children (p=0.034), and both groups displayed more *synchrony* than SC children (p(bonf)=0.037; >0.001, respectively); however, by adulthood, no difference between FT and KC groups was found (mean difference (MD)=−0.01, SE=0.12, p>0.05), BF_01_=3.52) and both displayed greater *synchrony* than SC participants (p=0.001). These findings show, for the first time, the long-term impact of maternal-newborn bodily contact on the mother-child relationship from infancy to adulthood.

**Fig. 2.**
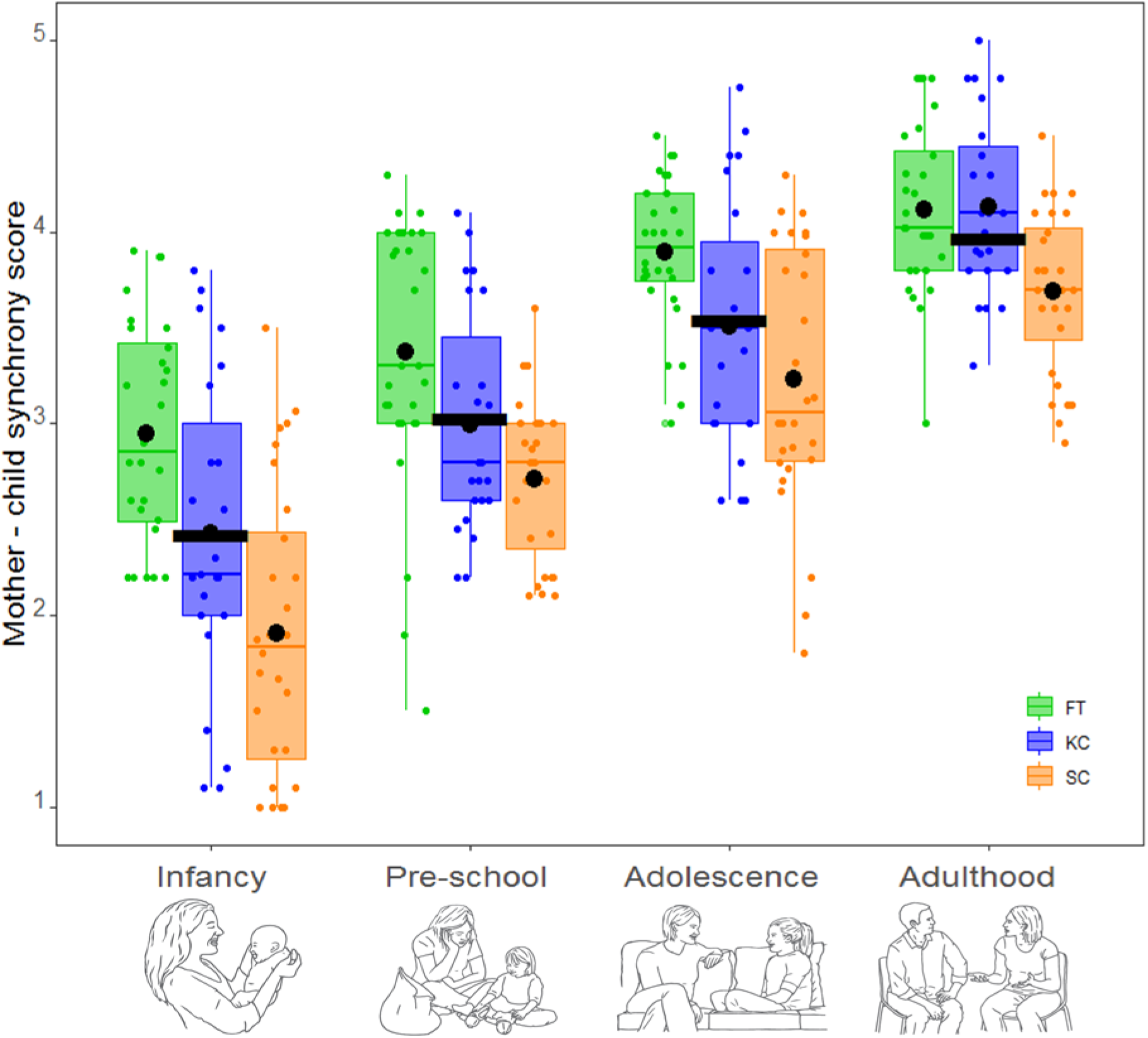
Mother-child synchrony from infancy to adulthood. Individual scores are marked by dots of respective color. Mean synchrony levels across participants are marked by black horizontal lines. Group means are marked by black circle. FT-Full-Term, KC-Kangaroo Care, SC-Standard Care

Next, we turned to assessing the neural systems implicated in empathy to distinct emotions by employing a validated paradigm ^13^ that exposed participants to protagonists experiencing joy, distress, or sadness in social contexts. Participants were instructed to “put themselves in the shoes” of the protagonists, to intentionally empathize with their feelings, and to imagine how they would feel in similar conditions (Fig. 1B and Supplementary Materials, SM).

Independent definition of regions of interest (ROIs) was ensured by dividing the dataset into two separate experiments. In Experiment 1, data from 18 subjects (six from each group: FT, KC, SC) were randomly selected for use as a functional localizer. Univariate whole brain mapping of emotional empathy (all emotions>scrambled image) revealed activated regions from the parietal-occipito border of the temporal lobe, across the superior temporal sulcus (STS) to TP, as well as prefrontal and limbic regions (amygdala, parahippocampal gyrus) and visual precuneus (Fig. 3A, Supplementary Table S1 for full details). Based on Experiment 1, we defined ROIs including the dorsomedial prefrontal cortex (DMPFC), ventromedial prefrontal cortex (VMPFC), precuneus, bilateral amygdala, bilateral (posterior) parahippocampal gyrus, bilateral inferior parietal cortex, bilateral STS, bilateral TP, and bilateral Insula, to be tested using RSA in the second, independent dataset (N=79). For visual demonstration of consistency, identical maps showing similar activation pattern in the two experiments appear in Fig. 3B and Fig. S1.

**Fig. 3.**
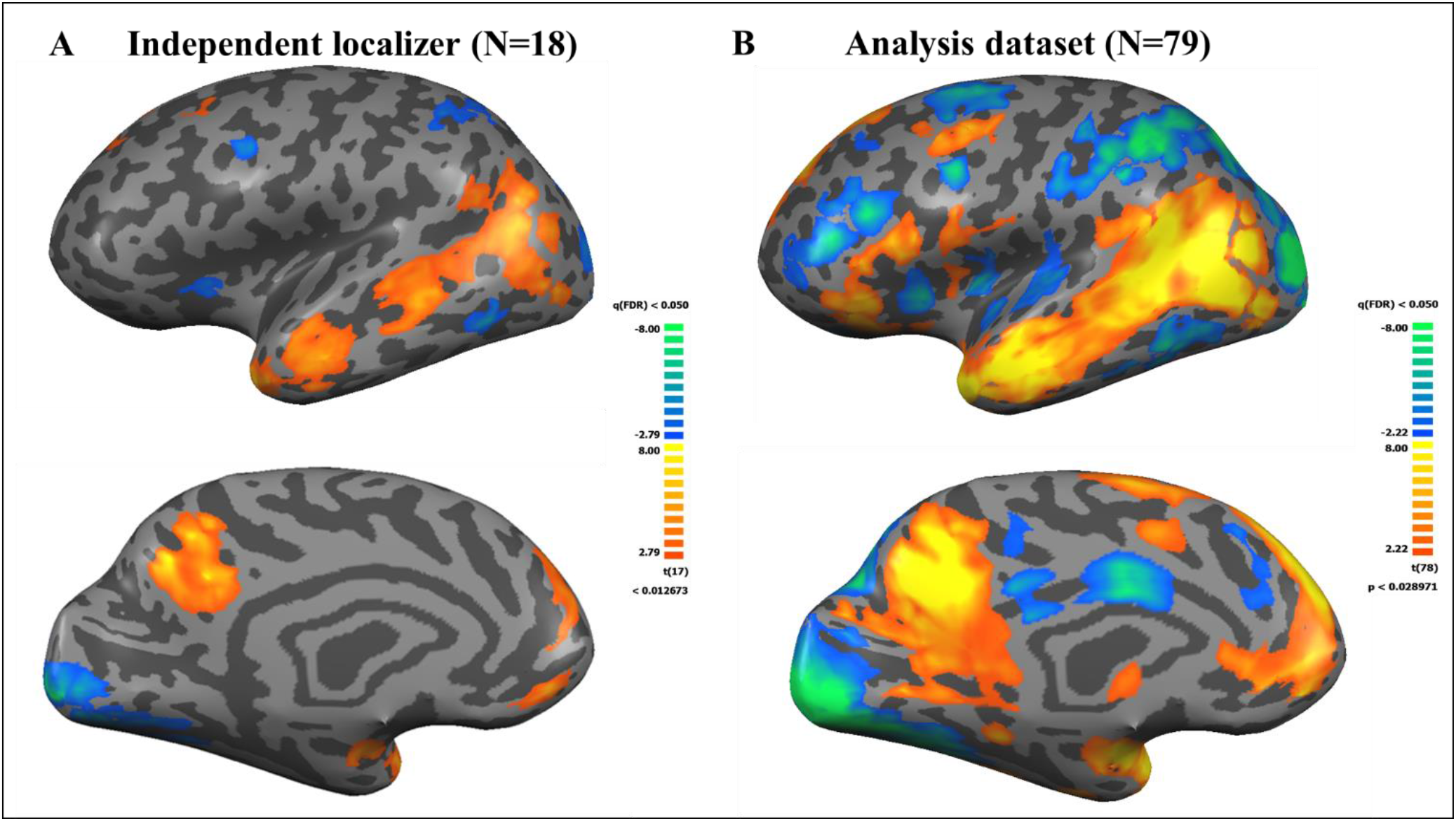
Whole brain maps of empathic accuracy. **A**: *First sample.* 18 subjects (6 from each group, FT, KC, SC) were randomly chosen for functional localizer. Presented contrast: emotion>scrambled image and group means used for ROI definition. **B**: *Second sample*: Similar contrast presented for analysis dataset (N=79). Note resemblance of activation patterns across samples. All maps are FDR corrected at q<0.05. Note: Images presenting left hemisphere. For right hemisphere see SM, fig S1.

Next, the representational geometry of empathy to different emotions was assessed. Using RSA (CoSMoMVPA toolbox ^17^) we examined the similarity of neural representations among emotions in each ROI separately in Experiment 2 (N=79, FT=28, KC=23, SC=28). For each ROI, we correlated the beta values of each voxel between each pair of emotions (joy-sadness, joy-distress, sadness-distress). Dissimilarity was defined as the correlation distance (1 - Pearson correlation) between each two conditions, calculated for each subject. The dissimilarity vectors for all 79 subjects were averaged ^18^ resulting in 3*3 dissimilarity matrix (DSM – see supplementary methods for full details). High dissimilarity indicates differential patterns of activation within a given ROI, denoting sensitivity of this region to different emotions ^14^.

RSA analysis revealed that the representational geometry of empathy to distinct emotions differed significantly between ROIs (*F_(5.2,400)_=77.17, p<0.001, η^2^_p_ =0.504*, GG sphericity correction) and between emotional pairs (*F*_*(1.6,1*27)_=9.41, p<0.001, η^2^_p_ =0.11); with joy-distress dissimilarity levels significantly higher than joy-sadness and sadness-distress (p<0.001). Significant ROI-dissimilarity-pair interaction effect emerged (*F_(8.6,660)_*=4.115, p<0.001, η^2^_p_=0.05), supporting our hypothesis that certain brain regions display specific responses to different emotions while others respond similarly across emotions. Moreover, we found a consistent gradient of dissimilarity across emotional pairs, indicating that regions exhibit similar levels of dissimilarity across all emotions. The amygdala and TP showed the highest dissimilarity, with insula displaying slightly lower but still high dissimilarity. The VMPFC, DMPFC, STS and parahippocampal gyrus exhibited medium-level dissimilarity, and the inferior parietal cortex and precuneus showed minimal dissimilarity (Fig 4; Supplementary table S2; multi-dimensional secondary (MDS) analysis in supplementary Fig S2). No significant group differences emerged in dissimilarity levels across ROIs (p=0.542) and no interaction effect for ROI-group and dissimilarity-group (p=0.379; 0.590 respectively).

**Fig. 4.**
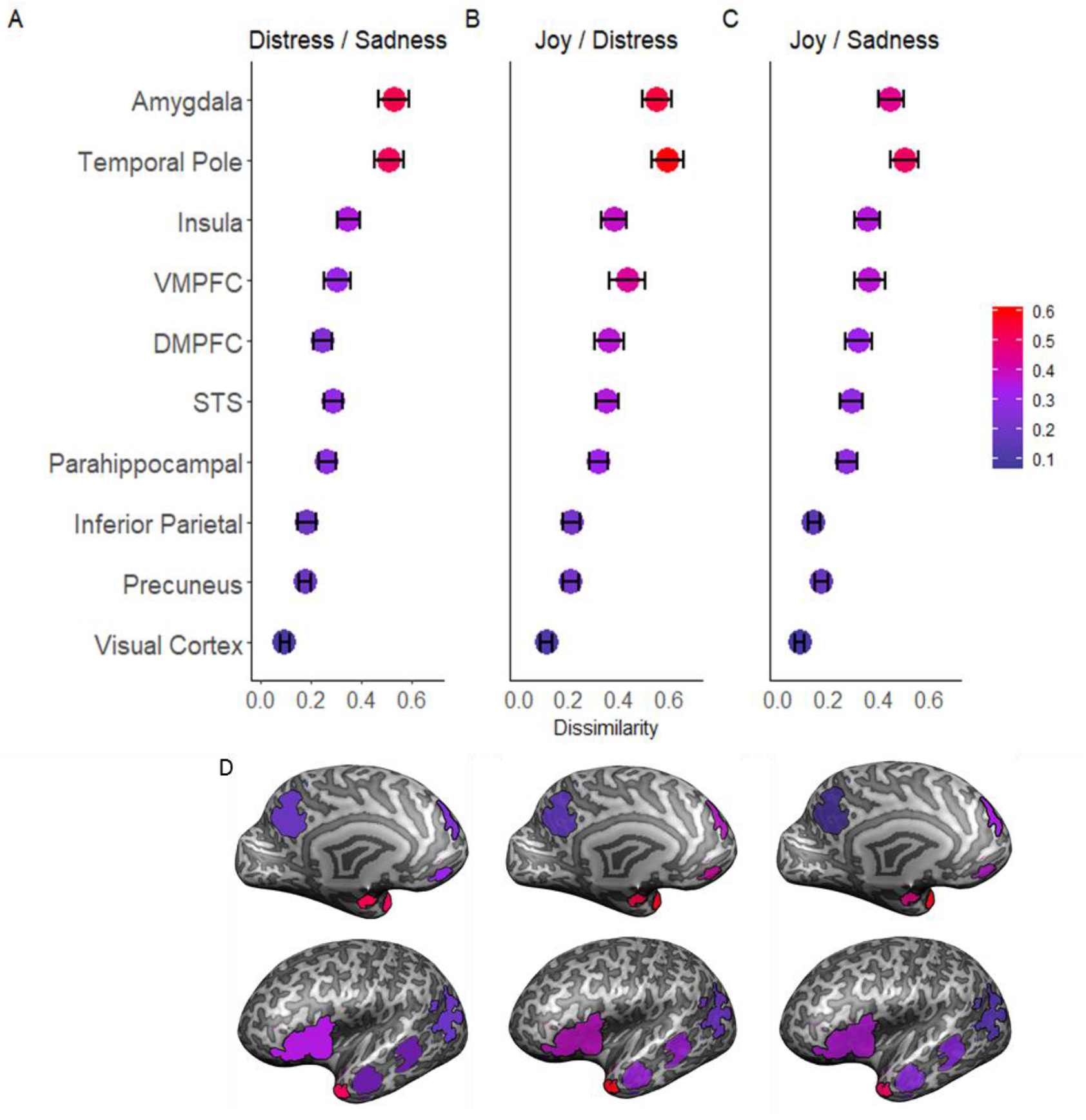
Representational similarity analysis across ROIs: Representational similarity was measured subject by subject by calculating correlation distance (1 minus Pearson correlation between two vectors of beta values of emotional pair) resulting in three dissimilarity pairs: (**A)** joy-distress, (**B**) joy-sadness, (**C**) distress-sadness for each ROI. Colour dots show mean dissimilarity levels for each ROI; error bars represent 95% confidence intervals. Similar pattern of dissimilarity are observed across emotional pairs. TP and amygdala show highest dissimilarity. Visual cortex (not ROI) presented for comparison of a low-level sensory region. (**D**) Actual ROIs, superimposed on an inflated brain. Color marks the dissimilarity levels, according to the color code of A-C.

We next moved to test our key hypothesis; the impact of mother-child *synchrony* across development (averaged from four time-points, see SM table S3) and the neural representation of empathy to others’ distinct emotions. We selected the three ROIs which were most sensitive to different emotions, areas for which dissimilarity levels for all three emotional pairs was above the median across all ROIs (see SM, table S4). The amygdala, TP, and insula were thus identified as emotion-sensitive regions based on their highest dissimilarity levels and were averaged into a *dissimilarity* score for each emotional pair.

Following, we tested the associations between *synchrony* and *dissimilarity* utilizing regression analysis with bootstrapping. We examined how *group* and *synchrony* predicted neural *dissimilarity* for each emotional pair. Results indicated that *synchrony* impacted dissimilarity for the joy-distress and joy-sadness pairs (bootstrapped *p*s<0.05), but not for the sadness-distress pair (p=0.13) (SM Table S5). These results support our key hypothesis, that mother-child synchrony experienced across development tunes the neural empathic response to distinct emotions, particularly valance-distinct emotions. However, a group-by-synchrony interaction followed by slope analysis indicated that while synchrony impacted *dissimilarity* in the FT and KC groups, contrary to our prediction, no such effect was found for the SC group (Figure 5A-C, SM table S5, S6; See Figure S3 and Table S8 for analyses by ROIs).

**Fig. 5.**
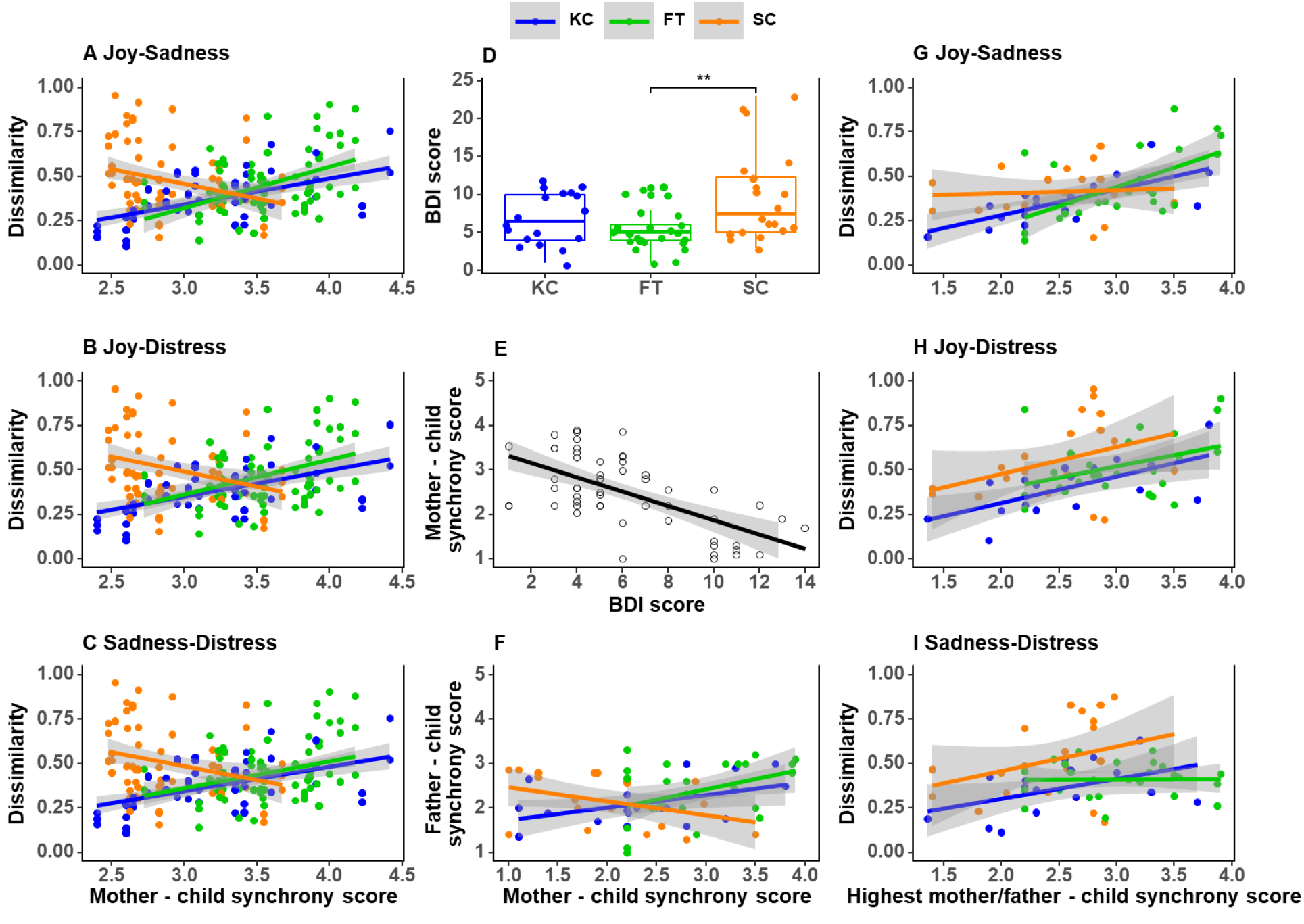
Impact of parent-child synchrony on adults’ neural empathic response. Regression analysis with bootstrap (5000 samples) to estimate the effects of mother-child synchrony across development on adults’ neural empathic accuracy in the three study groups for joy-distress (**A**), joy sadness (**B**) and sadness-distress (**C**). (**D**) Maternal depressive symptoms in infancy. (**E**) Correlation of maternal depressive symptoms and mother-infant synchrony (**F**) Correlations between mother-infant and father-infant synchrony in the three study groups. Regression analysis estimating the effects of highest parent-infant synchrony (mother or father) on adults’ neural empathic accuracy for the three groups: **(G)** joy-distress, (**H**) joy-sadness (**I**) and sadness distress

We therefore turned to examine the surprising lack of association between mother-child *synchrony* and neural dissimilarity in the SC group. If we consider parent-child synchrony as the core mechanism by which the human brain is tuned to empathic accuracy, why was this relationship absent in SC children and what was the compensatory mechanism sustaining neural empathy in these children? Data presented in Figure 2 indicated that the SC group had significantly lower *synchrony* compared to other groups. Prior work indicates that maternal depressive symptoms are consistently associated with reduced mother-infant synchrony ^19^. Indeed, our data from this cohort reveal significantly higher maternal depressive symptoms in the SC compared to the FT group (*F_(2,60)_*=5.77, *p*<0.01, *η^2^_p_* =0.16; Figure 5D) and maternal depressive symptoms correlated with lower synchrony (Figure 5E), suggesting that lower *synchrony* was driven by depressive symptoms. Thus, we postulated that the maturation of the neural empathic response was probably driven by synchronous interactions from another source.

Among bi-parental mammals, fathering is facultative and occurs in the context of maternal care where mother and father coordinate their caregiving to maximize offspring survival benefits ^20^. We previously found such co-parental complementary mechanisms in humans; when mothers are depressed, fathers step up and engage in greater synchrony and the increased father-infant synchrony buffers the long-term effects of maternal depression on child psychopathology ^21^. In such cases, the typical similarity between mother-infant and father-infant synchrony is not observed due to the complementary nature of their caregiving. Indeed, our data revealed positive correlation between mother-infant and father-infant synchrony in the FT and KC groups, but negative (marginal) correlation (*p*=0.07) in the SC group, suggesting complementary mechanism (Figure 5F; table S9).

We thus examined if complementary mechanism based on father-infant synchrony supported the longitudinal development of *dissimilarity* in the SC group. Father-child synchrony data was available only for infancy. We therefore used bootstrapped regressions to test whether the *highest level of synchrony* infants received from mother *or* father, predicted neural *dissimilarity* twenty years later (figure 5G-I and supplementary table S10). Results indeed indicated significant effects of *highest level of synchrony* for all three emotional pairs (*p*<0.01). Slope analyses showed that for the SC group, bootstrapped p-values were significant in the joy-distress and sadness-distress pairs but not for the joy-sadness (Table S11). These findings further confirm our main hypothesis that parent-child synchrony is the core mechanism for tuning the human brain to emotionally-accurate empathic response. Moreover, we demonstrate for the first time the existence of a complementary bi-parental mechanism in maturation of the human brain, where mother and father jointly modulate their caregiving according to family-level conditions and the provisions human infants receive from *either* parent impact the social brain in adulthood.

## Discussion

While findings in animal models provide evidence for the lifelong effects of mothering on the brain basis of sociality, ours is the first study to follow human infants across their lengthy maturation from birth to adulthood and to demonstrate the long-term impact of maternal proximity and synchronous caregiving on the adult brain. We capitalized on a unique “natural experiment”; the provision of maternal-newborn bodily contact during period of separation, followed by twenty years of repeated measurement of mother-child synchrony, and culminating in imaging the neural empathic response in adulthood. This enabled us to track how variations in initial conditions alter maternal caregiving, which, in turn, sharpen the brain’s capacity to empathize with others’ distinct emotions. Alterations in initial conditions resulted not only from maternal-infant separation but also from its ensuing effects on mother’s depressed mood and the father’s ability to step up and provide synchronous caregiving, addressing how *family-level* mechanisms by which bi-parental mammals coordinate caregiving according to changing ecological conditions, impact maturation of the human social brain. These findings support our main hypothesis: parent-child synchrony marks a core mechanism by which the human brain is tuned to the social world. Overall, our study is first to show how early caregiving patterns stabilize across lengthy periods and, over time, alter brain functioning, delineating how human relationships become brain.

Several key findings emerged from our twenty-year effort that may shed new light on three domains. First, we show that early relationships, with mother *or* father, longitudinally impact the social brain; second, we describe the neural structures presenting distinct representational geometries for empathy to different emotions; finally, we specify the neural network constituting the brain’s global empathic response to others’ emotions.

As expected, synchrony developed over time in all children and while infants can only momentarily engage in give-and-receive exchanges, adults can sustain long episodes of reciprocity, intimacy, and perspective-taking. Consistent with dynamic systems’ theory ^8^, synchrony was highly sensitive to initial conditions and adults born prematurely, especially those whose mothers reported high depressive symptoms did not show catchup and displayed consistently lower synchrony compared to their peers.

Maternal-newborn contact altered these initial conditions and our findings are first to demonstrate the pervasive effects of a neonatal touch-based intervention on the adult human. Touch triggered a different developmental trajectory, which, despite the fact that contact children showed mid-level synchrony across childhood, they displayed full catchup by adulthood. Following the pubertal transition, these children were able to engage in full-blown adult-adult synchrony, regardless of their high-risk beginning. Furthermore, the slope analysis demonstrated that synchrony experienced across development shaped the neural empathic response in this group. Possibly, one function of the maternal-newborn bodily contact is to provide a bridge from prenatal life, when maternal physiological systems are tuned to the fetal growth needs, to postnatal social life, when moments of interaction synchrony externally-regulate the infant’s heart rhythms, hormonal response, and brain oscillations and tune them to social life ^1,9^. When contact is eliminated during the post-partum period, the smooth transition of the maternal external-regulatory function breaks and the possibility of recovery may be limited.

Initial conditions involved not only maternal-infant attachment but also co-parental dynamics. The evolutionary pressures leading to social monogamy are unclear, but once monogamous mating systems stabilized they led to the emergence of paternal care and complex social behaviors. Bi-parental caregiving marks the first extension of the mother-infant bond to the larger social group and obligates parents to coordinate their effort to increase infant survival benefits, exposing young to multiple coordinated relationships within the family unit. Here we show for the first time the long-term impact of paternal caregiving on the human brain, indicating that such complementary mechanisms are as advantageous to human infants as they are to other bi-parental mammals ^20^.

Our study, employing multivariate pattern analyses in the empathy network, pinpointed structures that show distinct geometries to empathy for different emotions from regions indifferent to emotional content. Neural empathic accuracy was most prominent in the TP, amygdala, and insula. Anatomically, these regions are highly interconnected through the uncinate fasciculus, originating in the TP and forming the temporo-amygdala-orbitofrontal network ^22^. Through this interconnectivity, TP receives projections from the insula and projects to the amygdala, corroborating its role as paralimbic region ^23^.

Functionally, all three regions activate during empathy tasks (Fig. 3 & ^6^ and their activity is associated with emotional intensity ^24^. Here, as expected, the amygdala exhibited differential responses to empathy for different emotions, consistent with its documented role in emotional and social processing ^25^. The insula showed similar pattern of dissimilarity, consistent with its role in parent-infant synchrony ^26^, intersubjectivity, and empathic response to physical pain and emotional distress ^27,28^. The insula has been implicated in parental care and empathy across species ^29^ and has well-documented role integrating interoceptive and exteroceptive signals critical for the formation of a sense self ^30^, essential for empathic and affective responses ^31^.

Imaging studies show that the TP activates in Theory of Mind (ToM) and mentalizing tasks ^32^. Our findings are the first to show that the TP encodes distinct representational patterns for empathy to different emotions. TP lesioned old-world monkeys fail to recognize social signals and TP damage in humans is linked with impaired emotion recognition, poor social cognition, and inability to integrate facial, visual, and postural cues to specific persons. TP is involved in integrating emotionally-relevant multimodal sensory information into higher-order concepts that gauge the social-emotional significance of group members, critical for managing life in changing social environments. In monkeys, TP volumetric measures correlate with size of social group and in humans, with social network size ^33^.

Our findings extend these data by highlighting TP’s role in the online integration of perceptual, interoceptive, and emotional-affective stimuli ^34^. We suggest a putative model for insular-amygdalar-TP function in the empathic process; visceral bodily signals of interoceptive state originating in the insular cortex integrate with externally-oriented affective information from the limbic system and TP regions to enable modulation of the empathic stance in light of changing social contexts and arousal dynamics.

In addition to regions displaying emotion-specific representational geometries, results revealed a network of regions implicated in global affect sharing with no affect-specificity. Widespread regions of the parietal and frontal cortices were activated by the task. In the parietal cortex both the precuneus and inferior parietal cortex, extending to the TPJ and STS were strongly associated with the vicarious stance, but showed no emotion-specific modulations. These regions, components of the default mode network (DMN), have been associated with emotion processing ^35^, TOM, ^36^, and self-referential mentalizing ^37^ but not with discrete emotion representations in empathy contexts. Our results extend experimental ^38^ and meta-analytic ^39^ investigations on empathy, which typically focused on pain or distress, to distinguish the role of emotion-sensitive (TP, amygdala & Insula) from mentalizing regions (precuneus, STS, mPFC) in emotional empathy. Findings of the MDS analysis (figure S2) revealed that emotion-sensitive and mentalizing-general regions clustered together, suggesting distinct sub-systems sustaining aspects of empathy based on representational characteristics.

Understanding developmental continuity and change in humans- whose brain matures across three decades, involves substantial plasticity, and enables significant recovery following initial insult- is the focus of conceptual and empirical interest, particularly factors that tilt infants toward resilient trajectories. Integrating experimental design related to maternal-newborn contact and following infants for two decades, we found evidence for the two mechanisms proposed to underlie developmental continuity: step-by-step progress where minor alterations gradually impact outcome through repeated iterations, and a “sensitive period” perspective in which the system’s initial conditions directly impact long-term outcomes ^8^. Synchrony functioned as a core mechanism underpinning social development both by initiating a step-by-step trajectory that incorporates gradually-acquired socio-cognitive abilities and via direct impact of maternal or paternal synchrony on the adult brain. Our results indicate that the human social brain, like that of other mammals, is sensitive to features critical for human social life; rapid assessment of others’ feelings, accuracy of emotional detection, and interpretation of others’ mental states by assuming the vicarious stance. Our study, therefore, sharpens questions for future research and underscores the field of developmental social neuroscience as a potentially fruitful perspective on the human brain’s capacity for social living.

## Materials and Methods

### Participants

The study included 137 twenty-year old young adults (M=19.76, SD = 2.01, Range = 16.83-25.6) who were part of a longitudinal study recruited at birth and followed at our lab from birth until young adulthood. Of the study participants, 79 were born neurologically-intact preterm and were, at the time of brain imaging, healthy, well-functioning, and had graduated from typical state high-schools. Fifty-eight participants were born at full-term without any medical complications (FT). All children were reared in two-parent families to mothers who were above 21 years at childbirth and with family income above the poverty cutoff. Of the preterm group, 35 received kangaroo care (KC) during the hospitalization period in the Neonatal Intensive Care Units (NICU) (see below) and 44 infants, matched for demographic and medical variables, received standard incubator care (SC). See table S11: Demographics.

### Longitudinal design

The study utilized a birth-cohort longitudinal design, and mother-child dyads in both the preterm and full-term cohorts were recruited at birth during the same period.

#### Premature cohort

The original premature cohort and their mothers was recruited in Neonatal Intensive Care Units (NICU) at birth from March 1996 to November 1999, prior to any available information on the Kangaroo Care intervention and any scientific evidence for its-long term benefits and this enabled our randomization protocol (for details see ^40^. Because the developmental benefits of the Kangaroo Care intervention are now readily available to the public and most Western NICUs encourage mothers to engage in KC contact, the recruitment of a similar cohort is no longer possible and our carefully-matched cohort provides a rare opportunity to study the long-term effects of maternal separation and structured contact on the developing brain in a human model.

The initial preterm sample at birth included 146 mothers and their premature infants born at an average birthweight of M=1270gr. (SD=343.49, Range=530-1720 gr.) and gestational age M=30.65 weeks (SD=2.76, Range=25-34 weeks). Of these, 73 infants received skin-to-skin contact (Kangaroo Care, KC) and 73 matched controls, case-matched for demographic and medical conditions, including gender, birth-weight, gestational age, medical risk quantified by the Clinical Risk Index for Babies, maternal and paternal age and education, maternal employment, and parity. Only neurologically intact infants who were raised in two-parent, low-risk families were included, in order to tease apart the effects of maternal separation and bodily contact from developmental risk associated with early life contextual stress. Medical exclusion criteria included intra-ventricular hemorrhage grades III or IV, perinatal asphyxia, and metabolic or genetic disease. Contextual exclusion criteria included single motherhood, teenage mothers, and family income below the poverty cutoff.

#### Kangaroo Care Intervention

We recruited only infants who required full incubator care and could not receive full maternal-infant bodily contact outside the kangaroo position due to concern of the loss of body heat. Thus, the intervention was targeted to a period when no full bodily contact was possible between mother and infant among controls (standard incubation care; SC). For the KC intervention, infants were taken out of incubators, undressed, and placed between the mother’s breasts. Infants remained attached to cardio-respiratory monitor and were observed by nurses who recorded exact time of contact. Mothers sat in a standard rocking chair and used a bedside screen for privacy. To be a part of the study, mothers committed to providing KC for at least 1 hour per-day for at least 14 consecutive days. Details of the recruitment, randomization, and procedures during the hospitalization period were described in numerous prior publications ^40–43^.

Premature mothers and infants in both the KC and SC groups were observed nine times following the intervention; at hospital discharge, at 3,6,12 and 24 months, at 5 and 10-12 years, and two times in young adulthood, once for a home-visit and within the next weeks for an MRI scan). Of the original sample, we were able to visit the homes of 93 participants (63.6% of the original birth cohort), of which 79 participants were scanned (84.9%). General attrition was mainly related to inability to locate families or families moving to distant locations (32, 21.9%). Attrition from the scanning session was due to inability to meet safety requirements (e.g., tattoos, metal implants, epilepsy, surgeries, metal dental appliances) (13 participants, 8.9%) or refusal to undergo magnetic imaging (8, 5.5%).

#### Full-term cohort

The original full term cohort included 135 subjects, of which 52 were scanned (38%). Families meeting inclusion criteria (healthy, dual-parent, mothers above poverty line) were recruited from birth records in Well-Baby clinics. Families were observed four time, at 4 months, 2 years, 5 years, 12 years, and in both a home visit and MRI in young adulthood identical to the preterm cohort and conducted at the same period.

We were not able to locate 33 subjects (24%), and 38 (28%) refused participation. 11 (8%) subjects were not scanned because they did not meet safety requirements, or because they refused to, one subject terminated the scan due to claustrophobia. Additional 6 FT subjects were separately recruited with no longitudinal data.

### Parent-child synchrony

Mother-child synchrony was observed four times; in infancy, preschool, early adolescence, and young adulthood. Father-child synchrony was observed once, during infancy. Interactions were videotaped for offline coding.

a. Infancy: In infancy (M= 4.26 months, SD =1.14), we videotaped mother-infant interaction in the home environment. Instructions were “play with your infant as you normally do” and five minutes of free natural play were videotaped. During this visit, a similar father-infant interaction was also videotaped and order of parent was counter-balanced.
b. Preschool: During a home visit (M=3.21 years, SD = 1.38), mothers and children played with a set of predetermined toys for seven minutes. Parents and children were given a box of toys which were used in previous research on symbolic play of children at this age and were selected to elicit the child’s creativity and imagination ^44,45^. Toys included: two dolls, bottle, blanket, tea set including 2 cups, 2 plates, sugar and milk pots, and a boiler pan, wallet, colored necklace, a pair of plastic sunglasses, a sponge, three work tools, two small cars, telephone, 2 pet animals and 2 wild animals, and a small tool set.
c. Adolescence: In early adolescence (M = 12.07 years, SD = 1.62), mother and child engaged in two discussion paradigms for seven minutes each, consistent with prior research ^40,46–48^ in the first, a positive-valance task, mother and child were asked to plan “the best day ever” to spend together, and in the second, a negative-valance task, mother and child discussed a typical conflict in their relationship. Each task lasted for 7 minutes.
d. Adulthood: In young adulthood (M=19.76 years, SD = 2.01), young adults and their mothers engaged in the same positive valance task, as in adolescence and an additional 7 minute support giving task, in which each partner shared with the other a topic that concerns him.

### Behavior coding

The Coding Interactive Behavior (CIB) system ^49^ was used to code the positive and conflict discussions. The CIB is a well-validated global rating system for coding social interactions that includes 45-52 codes (depending on age) aggregated into several theoretically-based composites. The CIB has been used in a large number of studies across 17 countries and yielding over 150 publications. The system shows construct and predictive validity, test-retest reliability, and sensitivity to cultural contexts, interacting partner, and multiple psychopathological conditions [for a review see ^50^]. Consistent with prior research, we used the *parent-child synchrony* CIB construct; ^46,47,51–53^, computed by averaging several CIB codes.

- In infancy the *parent-child synchrony* construct included the following codes; mothers adapt stimulation to the child’s arousal state and social signals; interactions are rhythmic and have fluency; dyads show give-and-receive reciprocity; interactions are relaxed and not tense; and parent and child express positive affect and engagement (alpha =.91).
- In preschool, in addition to the infancy codes, the following codes were added to the *parent-child synchrony* construct; mother and child expand each other’s communications and symbolic expression, interactions are creative and non-restricted (alpha = .89).
- In adolescence, in addition to the infancy and preschool codes the following codes were included in the *parent-child synchrony* construct; parent and child acknowledge each other’s perspective, and parent and child show empathy to the partner’s positions, difficulties, complaints, or suggestions (alpha = .85).
- In adulthood, the parent-child synchrony construct included all these codes in addition to diminished hostility, withdrawn position, or slighting innuendos (alpha = .82).

Father-infant synchrony was computed using the same codes as mother-infant synchrony (alpha = .88). Coding was computed by trained coders blind to all other information. Inter-rater reliability computed on 15% of the sample and interrater reliability averaged, intraclass r = .93 (range: .88 - .99).

Parent-child synchrony showed moderate individual-stability from infancy to adulthood between all four observations (see Table S3), attesting to the stability of the synchronous caregiving style across the entire period of child development.

### Depressive symptoms

Maternal depressive symptoms score was measured with Beck depression inventory ^54^ during the infancy home visit and at young adulthood.

### MRI data acquisition

Magnetic Resonance Imaging (MRI) data was collected using a 3T scanner (SIEMENS MAGNETOM Prisma syngo MR D13D, Erlangen, Germany) located at the Tel Aviv Sourasky Medical Center. Scanning was conducted with a 20-channel head coil for parallel imaging. Head motion was minimized by padding the head with cushions, and participants were asked to lie still during the scan.

High resolution anatomical T1 images were acquired using magnetization prepared rapid gradient echo (MPRAGE) sequence: TR=1860ms, TE=2.74ms, FoV=256mm, Voxel size=1×1×1mm, flip angle=8 degrees.

Following, functional images were acquired using EPI gradient echo sequence. TR=2500ms, TE=30ms, 42 slices, slice thickness=3.5mm, FOV=220mm, Voxel size=3.2×2.3×3.5mm, flip angle=82 degrees. In total 393 volumes were acquired over the course of the empathy paradigm.

Visual stimuli were displayed to subjects inside the scanner, using a projector (Epson PowerLite 74C, resolution = 1024 × 768), and were back-projected onto a screen mounted above subjects’ heads, and seen by the subjects via an angled mirror. The stimuli were delivered using “Presentation” software (www.neurobs.com).

Before participating, participants signed an informed consent according to protocols approved by ethics committee of the Tel-Aviv Sourasky Medical Center. Subjects received a gift certificate of 450 NIS (~100USD) for their participation. The study was approved by Bar Ilan university’s IRB and by the Helsinki committee of the Sourasky medical center, Tel Aviv. Ethical approval no. 0084-15-TLV.

### Empathy paradigm

Empathy paradigm was adapted from Morelli et al. ^13^. During the fMRI scan, participants saw images of targets experiencing joy, sadness or distress-eliciting emotional events embedded in a social context.

The joy context-dependent experiences included images of protagonists looking joyful along with contextual information (for instance, “this person has just won the lottery”). The sadness context-dependent experiences included images of protagonists looking sad with contextual information (“this person just lost an important sport competition”). The distress context-dependent experiences comprised images of protagonists looking stressed and anxious with contextual information of uncertainty (“this person might lose his job”). In addition to emotional stimuli, a “neutral” condition included targets performing mundane non-emotional actions (“this person is ironing his shirts”). In order to capture sensory visual input alone, a “scrambled image” condition was included consisting of randomly shifted pixels of the photos presented in the other conditions. The instruction for this condition (to replace context) was “look carefully at the following images”.

Subjects were explicitly instructed to “put themselves in the shoes” of the protagonists, intentionally empathize with their feelings, and imagine how they would feel in similar situations. Overall, the session included 5 conditions (joy, sadness, distress, neutral, scrambled image), with each condition running for 5 blocks in a pseudo-random order ascertaining that 2 consecutive blocks are not of the same condition. The context (sentence) was presented for 2.5 seconds, followed by 4 photos presented for 5 seconds each (4*5=20 seconds). Between the blocks there was a fixation period of 10/12.5 seconds to avoid anticipation of the next stimulus. During fixation participants saw a white fixation cross in the middle of the screen over a black background (see figure 1B for details.

### Empathy paradigm validation

Prior to the fMRI experiment, stimuli were validated by independent raters (n = 11). Each rater was asked to rate the affective valence of each stimuli on a scale of 1 (very negative) to 7 (very positive). The valence for the “joy” condition stimuli was rated as positive (median=6.27, SD=0.64) and the valance for the “sadness” and “distress” conditions stimuli was rated as negative (“sadness”: median=2.36, SD=0.50; “distress”: median=3.09, SD=0.53) and the difference between categories was highly significant (p<0.001). Rating also considered the level of arousal for each stimuli on a scale of 1 (no arousal) to 7 (very high arousal). The “joy” condition stimuli were rated as high arousal (median =6.09 SD=0.83), whereas stimuli for the ‘distress’ and ‘sadness’ conditions were rated as significantly less arousing (median=4.72, SD=0.90; median=5.27, SD=0.90 respectively). Following, a one sample T-test showed that ‘distress’ stimuli median arousal ratings are greater than 3.5 (=medium-high arousal, t=4.5, p<0.001). However, for the ‘sad’ stimuli arousal ratings, one sample T-test indicated that the median arousal ratings were not significantly lower than 3.5 (=low arousal, t=6.5, p=1) suggesting that the sad stimuli were mildly emotionally arousing. Context sentences of the different emotional conditions did not differ in word count, nor in number of characters (M=7.86, SD=1.24 words, M =34.80, SD=6.08 no. of characters; F(2,12) =2.81, p=0.10; F(2,12)=0.23, p=0.794). Additionally, stimuli were examined for physical parameters, including complexity, contrast, and brightness, and showed no statistically significant difference on any of these parameters between conditions (F(2,57)=1.43, p=0.24).

## Data Analysis

### Data preprocessing

Data preprocessing and data analysis was done by BrainVoyager QX software package 20.6 (Brain Innovation, Maastricht, The Netherlands) ^55^. The first 3 volumes, before signal stabilization, were discarded to allow for T1 equilibrium. Preprocessing of functional scans included 3D motion correction, slice scan time correction and spatial smoothing by a full width at half maximum (FWHM) 4-mm Gaussian kernel. The functional images were then superimposed on 2D anatomical images and incorporated into the 3D datasets through trilinear interpolation. The complete dataset was transformed into Talairach space ^56^ using sinc interpolation. We used 4 mm smoothed data for whole brain maps. RSA was done using unsmoothed data.

### ROIs definition and functional localizer

We used a separate group of subjects to define our regions of interest (ROI) and validated these ROIs on the larger group of participants. A group of 18 subjects, 6 from each group, were used for ROI definition by a functional localizer and full details on the ROIs are presented in Table S3. The mean age of these 18 subjects was 19.44 years old (SD=1.79), 72% males, 94% right-handed. The subjects’ brain imaging data of the empathy paradigm was used for a multi subject GLM, using separate subject predictors. Based on the contrast of all emotions (joy, sadness, distress) > scrambled image, FDR corrected, activations were used as a functional localizer for ROI definition. Insular cortex alone was defined using a rectangle bounding box, as it was not clearly activated and was a region of interest based on current literature ^57^.

### Representational similarity analysis

Representational similarity analysis was done using MATLAB R2018a (MathWorks Inc.) with NeuroElf (J. Weber, http://neuroelf.net) and CoSMoMVPA toolbox ^17^. For each subject of the 79 ‘data analysis’ group, in each ROI, beta values per voxel for each condition (=empathy to a certain emotion, sadness, joy, distress) were extracted, resulting in 3 vectors (each vector is 1*number of voxels). Then, the correlation distance (=1 minus Pearson correlation) between every two conditions was calculated, for each subject. The dissimilarity vectors for all 79 subjects were averaged, resulting in 3*3 dissimilarity matrix (DSM). Following, in order to compare between different ROIs, a multi-dimensional secondary level was used, in which the correlation distances of each ROI DSM were projected into a two dimensional space ^58^. In addition, to define ‘high dissimilarity ROIs’, the correlation distance was transformed into percentage. As the values of dissimilarity range between 0 to 2, the percentage was calculated as dissimilarity value/2*100. We then calculated the mean (15.77) and the median (13.90%) percentage of dissimilarity were calculated across all ROIs. As the dissimilarity percentages are not normally distributed, the median (and not the mean) was preferred and “high dissimilarity ROIs” were defined as an ROI in which the median dissimilarity levels for all three emotional conditions are above the general median (>13.90%).

### fMRI analysis Exclusion criteria

#### (a) SC (Standard Care) group

In the SC group, 44 subjects were scanned. Of these, four were excluded due to excessive head movements (≥3mm), one was excluded due to a large cyst (~3cm), one was excluded due to technical problems with the scanner (only 385 of 393 dicoms were acquired), two were excluded due to failure in coregistration because of wrong positioning in the scanner, and two subjects were excluded due to head movement pattern, which affected coregistration.

#### (b) KC (Kangaroo Care) group

In the KC group, 35 subjects were scanned. Three were excluded due to excessive head movements (≥3mm), one was excluded because he fell asleep during the scan, and two were excluded due to deformed ventricles.

#### (c) FT (Full Term) group

In the FT group, 58 subjects were scanned. Three were excluded due to excessive head movements (≥3mm), 3 were excluded because of head movements, which affected coregistration, 6 subjects were excluded because functional volumes were upside down and coregistration failed, one subject had a technical problem in the scanner and not all dicoms were acquired, 2 subjects were misplaced in the scanner, 2 subjects were excluded based on activation maps, 1 subject had no activation in the visual cortex, 1 had only right hemisphere activation, and 1 failed to warp into Tal space. For a summary of excluded subjects by group, see table S13.

In total, 137 subjects were scanned: 79 preterm and 58 full-term. Of these, 103 subjects were analyzed (29 KC, 34 SC, 40 FT).

### Statistical Analysis

Statistical analysis was conducted using JASP (Version 0.9 for windows, JASP Team, 2018), SPSS (SPSS statistics V25, IBM Corp.) and in R version 3.5.3 ^59^. Bootstrapping were calculated using the packages “boot”, “parameters”, and personal functions ^60–62^.

## Acknowledgments

We are grateful to the participants and their families for their kind cooperation across the years.

## Author contributions

R.F. conceived and designed the longitudinal study and conducted the twenty-year follow-up. A.U.Y analyzed the data. R.S. supervised the analysis. A.U.Y. and S.W. ran the experiments. A.D. performed regressions and statistical analysis. O.S.R. contributed to fMRI paradigm validation. A.U.Y, R.S. and R.F wrote the paper. R.F and R.S. further contributed to the writing by reviewing and editing the manuscript.

## Data and materials availability

The data that support the findings of this study are available on request from the corresponding author, R.F. The data are not publicly available due to its nature – videos of interactions containing information that could compromise the privacy of research participants.

## Supplementary Materials

Figures S1-S3

Tables S1-S13

References (*1-26*)

