## Supplementary tables and figures for "Synchronous Caregiving from Birth to Adulthood Tunes Humans’ Social Brain"

**This file includes:**

Supplementary Text
Figs. S1 to S3
Tables S1 to S13

### Supplementary Text

*Preregistration*

An earlier version of this study was pre-registered on open science framework website on October 2018. However, as we have enlarged the scope of the study considerably adding additional subject groups and analysis we amended this pre-registration. The annulled pre-registration details and amendments can be viewed using the link: https://osf.io/wra8x/?view_only=31a0013b2a264cf38ca992bdc3cdc594

*Controlling for age effects*

To address the age differences between groups (see table S11, demographics), we confirmed that dissimilarity levels were unrelated to participants' age. A univariate ANOVA analysis, corrected for age and group showed that the Amygdala, TP and Insular dissimilarity levels of joy-sadness, joy-distress and distress-sadness are not significantly different between subjects (joy-sadness corrected $F(3,75)=2.184$, $p=0.097$;joy-distress $F(3,75)=2.389$, $p=0.075$;sadness-distress $F(3,75)=1.666$, $p=0.182$).

The development of synchrony was found to differ among groups ($F(2,76)=8.873$, $p<0.001$; see figure 2) and post-hoc analysis indicated that levels of synchrony in the FT and KC groups were higher compared to the SC group ($p=0.001$;$p=0.002$ respectively, bonferroni corrected). We examined a corrected model in which participants' age in young adulthood was used as a covariate. Results showed that the significant difference between groups remained when controlling for age ($F(3,75)=6.301$, $p=0.001$). The participant's age was not the source of this variability $F(1,75)=1.127$, $p=0.292$) neither for synchrony levels in young adulthood nor for the averaged synchrony score across time (corrected $F(3,75)=13.006$, $p<0.001$; age at young adulthood corrected $F(1,75)=1.281$, $p=0.261$).

*TP laterality effect in RSA*

Despite the accumulating evidence for TP role as a key socio-emotional region, and following literature associating TP with language and semantic processing [1], we wanted to explore if TP's involvement may be due to its role in semantic and language processing in humans, as our task included a semantic component. We thus hypothesized that if the TP effect is of semantic origin it is likely that it will show a left lateralized bias, in line with typical lateralization of language systems.

To tackle this, a separate RM ANOVA analysis for left and right TP, was done, in order to examine if there is a difference in dissimilarity levels. The analysis revealed that there was no significant difference between right and left TP dissimilarity levels ($F_{(1,78)}$=2.27, p=0.13). There was a significant main effect for emotion pairs dissimilarity ($F_{(2,156)}$=4.38,p=0.014, $\eta^2_p$=0.053), similar to our bilateral ROI findings (fig. 4). Emotions-laterality interaction was also insignificant ($F_{(2,156)}$=0.541,p=0.58). Therefore, our findings indicate that the TP activation pattern is likely not based on semantic properties of the task, but on its social- emotional properties.

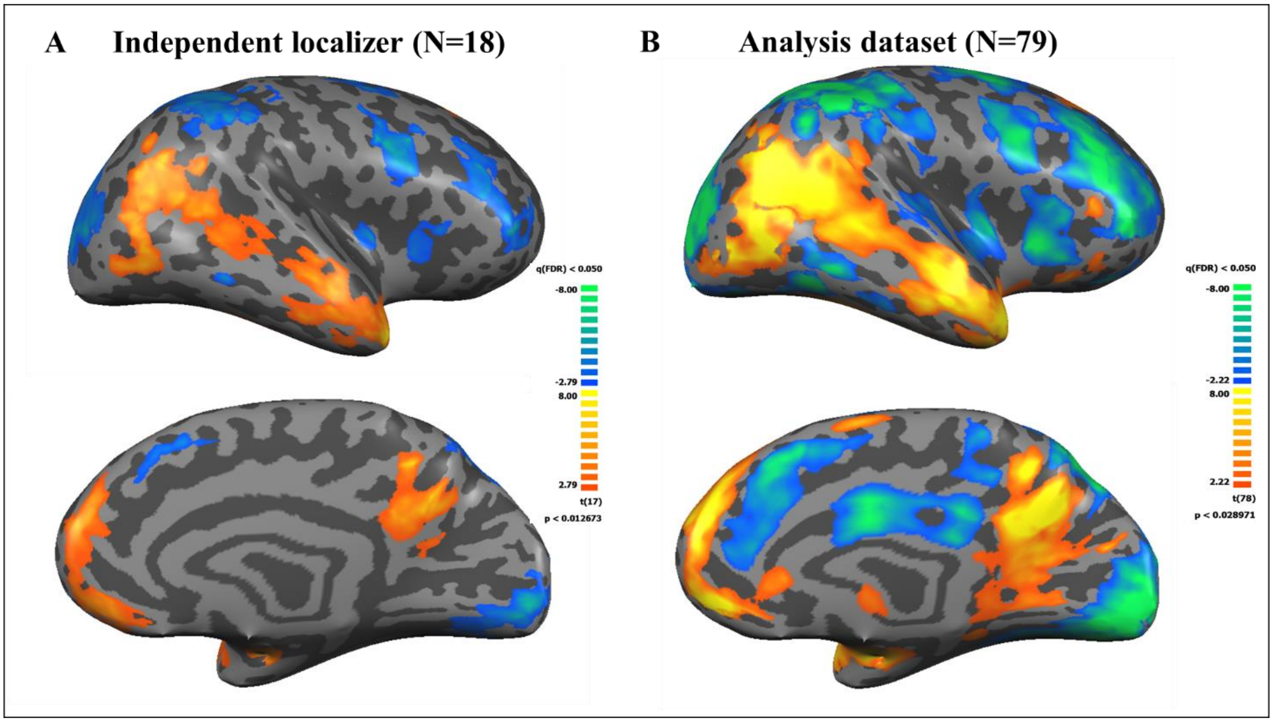


**Fig. S1.**

Whole brain maps of emotion. **A**: 18 subjects, 6 from each group (FT, KC and SC) were randomly chosen for functional localizer dataset. The presented emotion group map of 18 subjects was used for independent ROI definition, of the contrast emotion>scrambled image, presented on right hemisphere. **B**: same contrast, in the other 79 subjects (analysis dataset) - note the high degree resemblance of activation pattern between A and B. All maps are FDR corrected at q<0.05.

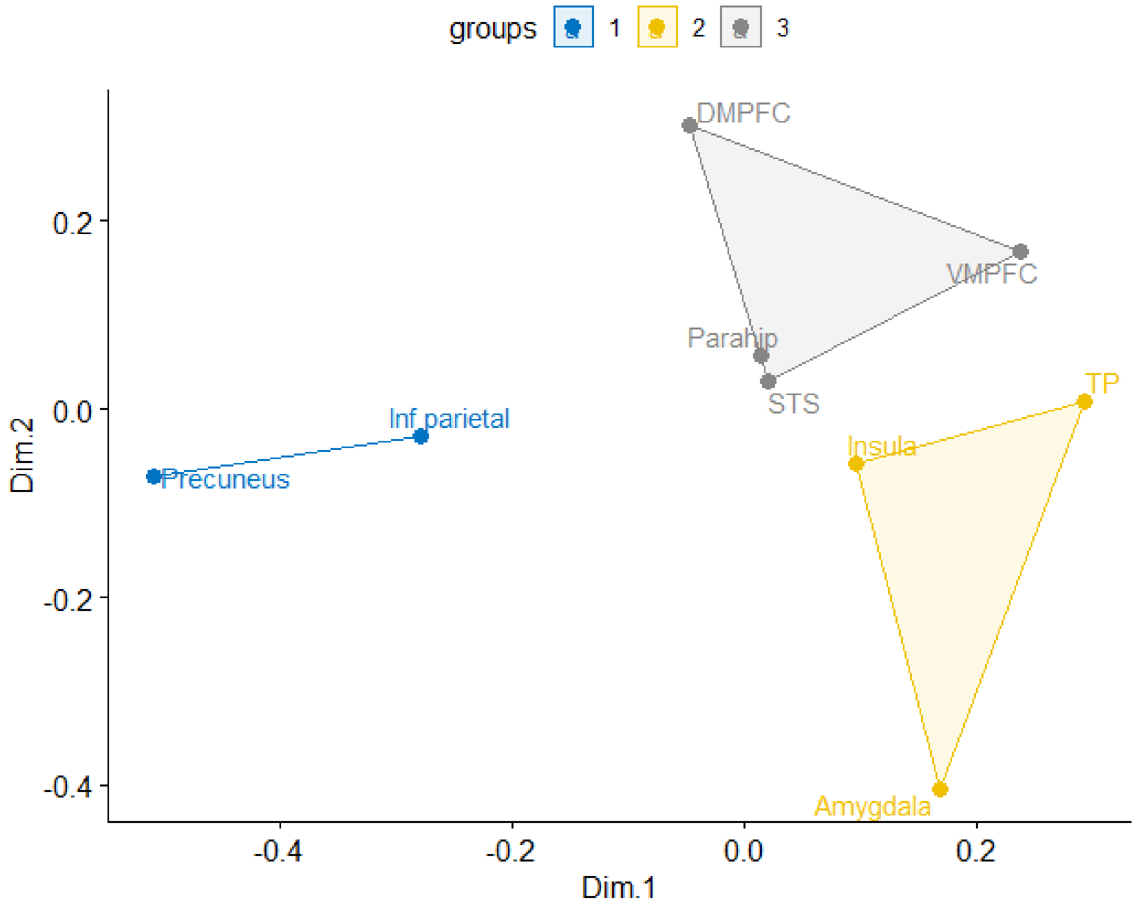


**Fig. S2.**

Multi-dimensional secondary levels analysis comparing between ROI's emotion specific representational geometries. Using k means clustering (k=3), we identified 3 groups based on the representation pattern of each region. Note the high dissimilarity levels regions – TP, insula and amygdala are clustered together (yellow), while the low dissimilarity regions inferior parietal cortex and precuneus cluster (blue) for a separate cluster.

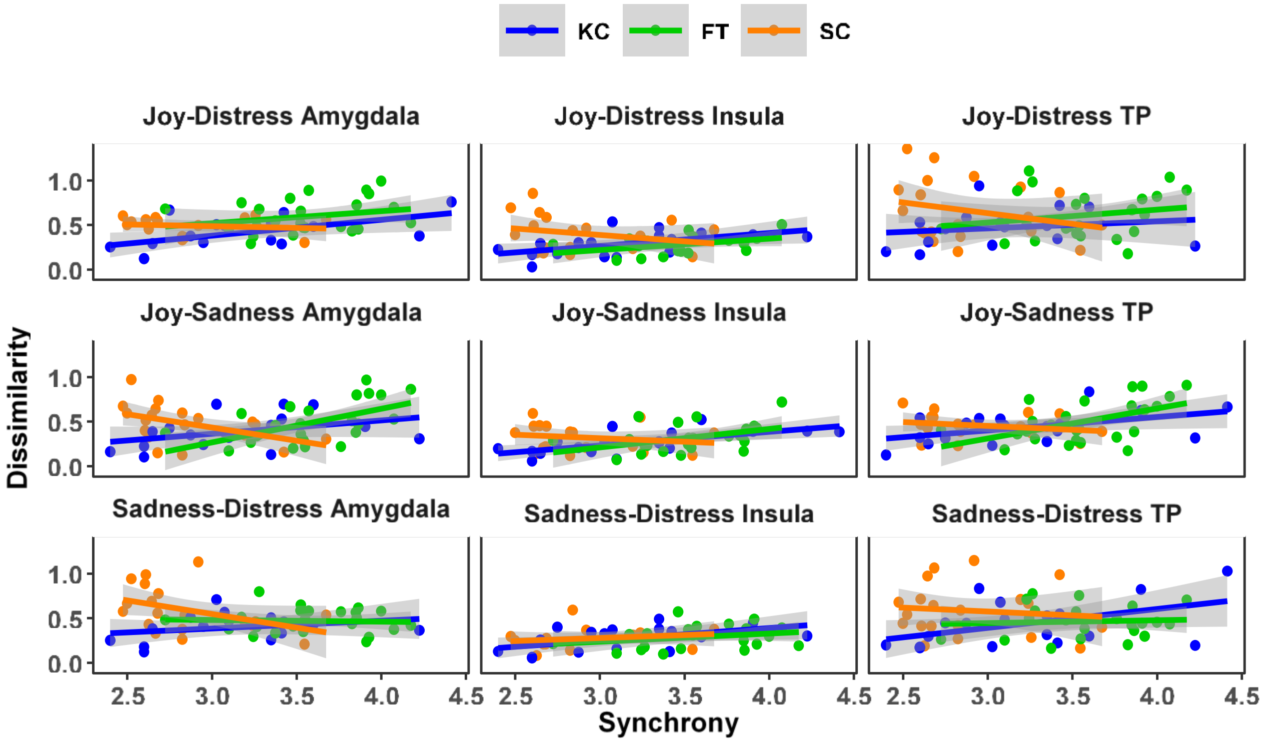


**Fig. S3.**

Bootstrapped regression for the relationship between behavioral synchrony and dissimilarity levels moderated by birth group

Note:

1. Bootstrapped with 5,000 samples.
2. As can be seen in Table S10., after bootstrap with 5,000 samples, p-values were significant for joy-distress in amygdala and insula for KC, for joy-sadness in amygdala in FT and SC, in insula in FT and KC, and for TP in FT and KC. Lastly, for sadness-distress in amygdala in SC, insula in KC, and for TP in KC

**Table S1.**

Regions active during emotional empathy

| **ROI** | | Mean X | Mean Y | Mean Z | STD X | STD Y | STD Z | K Cluster size | Peak X | Peak Y | Peak Z | t | p |
|---|---|---|---|---|---|---|---|---|---|---|---|---|---|
| Inferior Parietal cortex | R | 44.27 | -57.15 | 18.29 | 5.26 | 7.19 | 5.72 | 6179 | 45 | -61 | 25 | 7.480 | 0.000001 |
| | L | -44.92 | -62.45 | 20.62 | 4.83 | 5.19 | 6.15 | 4569 | -45 | -70 | 31 | 7.251 | 0.000001 |
| Precuneus | | -1.35 | -51.76 | 29.2 | 6.15 | 5.36 | 6.56 | 8327 | -6 | -55 | 34 | 9.471 | 0 |
| Temporal pole | R | 42.6 | 12.8 | -23.02 | 4.54 | 4.93 | 5.33 | 3570 | 42 | 17 | -26 | 9.331 | 0 |
| | L | -40.87 | 11.09 | -23.19 | 4.75 | 4.57 | 4.52 | 2983 | -42 | 17 | -23 | 8.080 | 0 |
| Superior Temporal sulcus | R | 50.58 | -23.75 | -2.42 | 4.89 | 19.61 | 14.61 | 7649 | 48 | -40 | 7 | 6.325 | 0.000008 |
| | L | -53.12 | -20.7 | -6.03 | 6.38 | 14.66 | 9.36 | 4239 | -54 | -40 | 1 | 7.196 | 0.000001 |
| Amygdala | R | 25.4 | -11.26 | -12.51 | 5.1 | 4.55 | 3.16 | 2565 | 30 | -4 | -14 | 9.789 | 0 |
| | L | -25.24 | -13.56 | -13.02 | 4.28 | 5.93 | 2.93 | 1796 | -33 | -7 | -14 | 7.869 | 0 |
| Parahippocampal gyrus | R | 36.62 | -36.75 | -16.09 | 2.02 | 4.54 | 3.04 | 541 | 36 | -40 | -20 | 5.399 | 0.000048 |
| | L | -36.14 | -39.44 | -16.14 | 1.61 | 1.74 | 2.07 | 214 | -36 | -40 | -17 | 5.504 | 0.000039 |
| VMPFC | | -0.76 | 46.11 | -3.71 | 4.31 | 5.15 | 3.8 | 2685 | -9 | 44 | -2 | 6.651 | 0.000004 |
| DMPFC | | -0.5 | 52.59 | 28.39 | 5.44 | 3.45 | 6.11 | 3223 | -3 | 53 | 34 | 5.904 | 0.000017 |
| Insula | R | 38.5 | 20.5 | 6 | 5.77 | 8.66 | 5.48 | 11400 | 36 | 17 | 4 | -4.698 | 0.000207 |
| | L | -38.5 | 20.5 | 6 | 5.77 | 8.66 | 5.48 | 11400 | -30 | 17 | 4 | -5.196 | 0.000073 |

**Table S2.**

Mean dissimilarity levels across ROIs

| ROI | Mean | Lower CI | Upper CI |
|---|---|---|---|
| | *Joy - Distress* | | |
| TP | 0.594 | 0.529 | 0.660 |
| Amygdala | 0.551 | 0.492 | 0.609 |
| VMPFC | 0.434 | 0.363 | 0.505 |
| Insula | 0.381 | 0.332 | 0.429 |
| DMPFC | 0.363 | 0.305 | 0.420 |
| STS | 0.352 | 0.307 | 0.396 |
| Parahippocampal gyrus | 0.320 | 0.282 | 0.357 |
| Inferior Parietal cortex | 0.212 | 0.179 | 0.245 |
| Precuneus | 0.208 | 0.176 | 0.241 |
| VC | 0.111 | 0.087 | 0.135 |
| | *Joy - Sadness* | | |
| | Mean | Lower CI | Upper CI |
| TP | 0.504 | 0.446 | 0.561 |
| Amygdala | 0.450 | 0.401 | 0.500 |
| VMPFC | 0.366 | 0.303 | 0.428 |
| Insula | 0.356 | 0.308 | 0.403 |
| DMPFC | 0.321 | 0.268 | 0.373 |
| STS | 0.294 | 0.248 | 0.339 |
| Parahippocampal gyrus | 0.275 | 0.235 | 0.315 |
| Precuneus | 0.174 | 0.148 | 0.200 |
| Inferior Parietal cortex | 0.143 | 0.119 | 0.166 |
| VC | 0.088 | 0.068 | 0.107 |
| | *Sadness-Distress* | | |
| | Mean | Lower CI | Upper CI |
| Amygdala | 0.533 | 0.473 | 0.592 |
| TP | 0.512 | 0.453 | 0.572 |
| Insula | 0.350 | 0.305 | 0.396 |
| VMPFC | 0.308 | 0.256 | 0.361 |
| STS | 0.290 | 0.252 | 0.328 |
| Parahippocampal gyrus | 0.266 | 0.231 | 0.301 |
| DMPFC | 0.247 | 0.211 | 0.283 |
| Inferior Parietal cortex | 0.185 | 0.149 | 0.221 |
| Precuneus | 0.179 | 0.155 | 0.203 |
| VC | 0.098 | 0.077 | 0.118 |

**Table S3.**

Synchrony correlations across child development

| | | Infancy | Preschool | Adolescence | Adulthood |
|---|---|---|---|---|---|
| **Infancy** | Pearson's r | — | | | |
| | p-value | — | | | |
| **Preschool** | Pearson's r | 0.584 | — | | |
| | p-value | **< .001** | — | | |
| **Adolescence** | Pearson's r | 0.529 | 0.496 | — | |
| | p-value | **< .001** | **< .001** | — | |
| **Adulthood** | Pearson's r | 0.492 | 0.398 | 0.462 | — |
| | p-value | **< .001** | **< .001** | **< .001** | — |

**Table S4.**

Mean and median percentage of dissimilarity across ROIs

The median dissimilarity levels across all ROIs was 13.901%. Regions were defined as "high dissimilarity" if their median dissimilarity percentage was higher than the general median for all three emotion pairs.

| ROI | Emotions | Median % dissimilarity | Mean % dissimilarity |
|---|---|---|---|
| Amygdala | Joy - Sadness | **21.837** | 21.706 |
| | Joy - Distress | **27.360** | 29.608 |
| | Sadness - Distress | **26.638** | 30.041 |
| TP | Joy - Sadness | **23.012** | 26.911 |
| | Joy - Distress | **24.830** | 30.593 |
| | Sadness - Distress | **26.961** | 29.270 |
| Insula | Joy - Sadness | **18.204** | 19.064 |
| | Joy - Distress | **18.666** | 20.590 |
| | Sadness - Distress | **15.452** | 18.789 |
| VMPFC | Joy - Sadness | **15.479** | 17.944 |
| | Joy - Distress | **20.004** | 23.553 |
| | Sadness - Distress | 10.128 | 15.303 |
| DMPFC | Joy - Sadness | 10.154 | 13.420 |
| | Joy - Distress | **16.002** | 17.909 |
| | Sadness - Distress | 10.489 | 11.216 |
| STS | Joy - Sadness | 10.311 | 14.396 |
| | Joy - Distress | 12.948 | 17.237 |
| | Sadness - Distress | 12.199 | 15.322 |
| Parahippocampal gyrus | Joy - Sadness | 12.326 | 14.299 |
| | Joy - Distress | 13.570 | 18.069 |
| | Sadness - Distress | 12.085 | 15.004 |
| Inferior parietal gyrus | Joy - Sadness | 5.124 | 6.856 |
| | Joy - Distress | 9.199 | 11.811 |
| | Sadness - Distress | 6.788 | 11.066 |
| Precuneus | Joy - Sadness | 6.152 | 8.222 |
| | Joy - Distress | 9.745 | 11.364 |
| | Sadness - Distress | 6.357 | 8.971 |

**Table S5.**

Bootstrapped regression for the relationship between behavioral synchrony and dissimilarity levels moderated by birth group

| | **Joy-Distress** | | | **Joy-Sadness** | | | **Sadness-Distress** | | |
|---|---|---|---|---|---|---|---|---|---|
| *Predictors* | *B* | *CI* | *P* | *B* | *CI* | *p* | *B* | *CI* | *p* |
| Group FT | -0.11 | -0.92 – 0.65 | 0.755 | -0.70 | -1.47 – 0.04 | 0.060 | 0.36 | -0.29 – 1.09 | 0.274 |
| Group SC | 1.19 | 0.33 – 2.05 | **0.009** | 1.01 | 0.49 – 1.58 | **0.001** | 0.95 | -0.03 – 1.86 | 0.056 |
| Synchrony | 0.16 | 0.02 – 0.29 | **0.029** | 0.15 | 0.07 – 0.29 | **<0.001** | 0.12 | -0.04 – 0.30 | 0.133 |
| Synchrony:Group FT | 0.05 | -0.18 – 0.29 | 0.645 | 0.20 | -0.02 – 0.42 | 0.074 | -0.10 | -0.33 – 0.09 | 0.312 |
| Synchrony:Group SC | -0.34 | -0.64 – -0.06 | **0.020** | -0.32 | -0.51 – -0.15 | **0.002** | -0.26 | -0.57 – 0.07 | 0.120 |
| $R^2$ / $R^2$ adjusted | 0.259 / 0.194 | | | 0.401 / 0.347 | | | 0.213 / 0.143 | | |

Note:

1. Confidence intervals (CI) and p-values are bootstrapped with 5,000 samples.
2. KC was used as a reference group.
3. Bs reflect the unstandardized regression coefficient

**Table S6.**

Bootstrapped for the slopes of the relationship between maternal behavioral synchrony and dissimilarity levels for each group

| | *Group* | *Est.* | *S.E.* | *t-value* | *CI - 2.5%* | *CI - 97.5%* | *P-value* |
|---|---|---|---|---|---|---|---|
| Joy - Sadness | FT | 0.352 | 0.074 | 4.728 | 0.206 | 0.498 | **0.000** |
| Joy - Sadness | KC | 0.147 | 0.056 | 2.628 | 0.037 | 0.257 | **0.009** |
| Joy-Sadness | SC | -0.148 | 0.080 | -1.848 | -0.304 | 0.009 | 0.065 |
| Joy - Distress | FT | 0.205 | 0.099 | 2.057 | 0.010 | 0.400 | **0.040** |
| Joy - Distress | KC | 0.156 | 0.075 | 2.087 | 0.009 | 0.303 | **0.037** |
| Joy - Distress | SC | -0.153 | 0.105 | -1.457 | -0.359 | 0.053 | 0.145 |
| Sadness - Distress | FT | 0.025 | 0.093 | 0.269 | -0.156 | 0.206 | 0.788 |
| Sadness - Distress | KC | 0.119 | 0.083 | 1.441 | -0.043 | 0.281 | 0.150 |
| Sadness - Distress | SC | -0.099 | 0.098 | -1.014 | -0.291 | 0.093 | 0.310 |

Note: Confidence intervals (CI) and p-values are bootstrapped with 5,000 samples.

**Table S7.**

Bootstrapped for the slopes of the relationship between maternal behavioral synchrony and dissimilarity levels for each birth group

| | *Group* | *Est.* | *S.E.* | *t-value* | *CI - 2.5%* | *CI - 97.5%* | *P-value* |
|---|---|---|---|---|---|---|---|
| Joy-Distress Amygdala | FT | 0.135 | 0.097 | 1.391 | -0.055 | 0.326 | 0.164 |
| Joy-Distress Amygdala | KC | 0.179 | 0.077 | 2.324 | 0.028 | 0.330 | **0.020** |
| Joy-Distress Amygdala | SC | 0.002 | 0.112 | 0.016 | -0.217 | 0.220 | 0.987 |
| Joy-Distress Insula | FT | 0.127 | 0.098 | 1.291 | -0.066 | 0.319 | 0.197 |
| Joy-Distress Insula | KC | 0.145 | 0.073 | 1.978 | 0.001 | 0.289 | **0.048** |
| Joy-Distress Insula | SC | -0.146 | 0.089 | -1.645 | -0.320 | 0.028 | 0.100 |
| Joy-Distress TP | FT | 0.145 | 0.158 | 0.916 | -0.165 | 0.454 | 0.360 |
| Joy-Distress TP | KC | 0.079 | 0.153 | 0.515 | -0.222 | 0.379 | 0.607 |
| Joy-Distress TP | SC | -0.225 | 0.165 | -1.361 | -0.549 | 0.099 | 0.174 |
| Joy-Sadness Amygdala | FT | 0.377 | 0.110 | 3.422 | 0.161 | 0.593 | **0.001** |
| Joy-Sadness Amygdala | KC | 0.151 | 0.098 | 1.536 | -0.042 | 0.343 | 0.125 |
| Joy-Sadness Amygdala | SC | -0.257 | 0.116 | -2.208 | -0.484 | -0.029 | **0.027** |
| Joy-Sadness Insula | FT | 0.207 | 0.087 | 2.374 | 0.036 | 0.378 | **0.018** |
| Joy-Sadness Insula | KC | 0.152 | 0.059 | 2.587 | 0.037 | 0.267 | **0.010** |
| Joy-Sadness Insula | SC | -0.089 | 0.088 | -1.017 | -0.261 | 0.083 | 0.309 |
| Joy-Sadness TP | FT | 0.338 | 0.099 | 3.413 | 0.144 | 0.531 | **0.001** |
| Joy-Sadness TP | KC | 0.151 | 0.075 | 2.025 | 0.005 | 0.297 | **0.043** |
| Joy-Sadness TP | SC | -0.079 | 0.111 | -0.709 | -0.296 | 0.139 | 0.478 |
| Sadness-Distress Amygdala | FT | -0.018 | 0.108 | -0.165 | -0.229 | 0.194 | 0.869 |
| Sadness-Distress Amygdala | KC | 0.087 | 0.107 | 0.818 | -0.122 | 0.296 | 0.413 |

| | | | | | | | |
|---|---|---|---|---|---|---|---|
| Sadness-Distress Amygdala | SC | -0.253 | 0.120 | -2.106 | -0.489 | -0.018 | **0.035** |
| Sadness-Distress Insula | FT | 0.092 | 0.068 | 1.355 | -0.041 | 0.225 | 0.175 |
| Sadness-Distress Insula | KC | 0.140 | 0.059 | 2.378 | 0.025 | 0.255 | **0.017** |
| Sadness-Distress Insula | SC | 0.084 | 0.076 | 1.113 | -0.064 | 0.232 | 0.266 |
| Sadness-Distress TP | FT | 0.033 | 0.142 | 0.231 | -0.246 | 0.312 | 0.817 |
| Sadness-Distress TP | KC | 0.213 | 0.104 | 2.043 | 0.009 | 0.417 | **0.041** |
| Sadness-Distress TP | SC | -0.053 | 0.146 | -0.362 | -0.339 | 0.233 | 0.718 |

Note: Confidence intervals (CI) and p-values are bootstrapped with 5,000 sample

**Table S8.**

Bootstrapped regression for the relationship between mother and father behavioral synchrony moderated by birth group

Submitted also as a separate file, tableS8.xlsx, due to size

| | **Joy-Distress Amygdala** | | | **Joy-Distress Insula** | | | **Joy-Distress TP** | | | **Joy-Sadness Amygdala** | | | **Joy-Sadness I** | |
|---|---|---|---|---|---|---|---|---|---|---|---|---|---|---|
| *Predictors* | *B* | *CI* | *p* | *B* | *CI* | *p* | *B* | *CI* | *p* | *B* | *CI* | *p* | *B* | *CI* |
| FT-KC | 0.26 | - 0.85 – 1.12 | 0.611 | 0.03 | - 0.62 – 0.59 | 0.932 | - 0.05 | - 1.38 – 1.52 | 0.945 | - 0.75 | - 1.81 – 0.28 | 0.147 | - 0.18 | - 1.06 – 0.59 |
| SC-KC | 0.77 | 0.05 – 1.51 | **0.035** | 1.02 | 0.21 – 1.92 | **0.010** | 1.22 | - 0.23 – 2.70 | 0.098 | 1.46 | 0.63 – 2.41 | **0.001** | 0.79 | 0.17 – 1.48 |
| Synchrony | 0.18 | 0.01 – 0.33 | **0.040** | 0.15 | 0.07 – 0.27 | **0.003** | 0.08 | - 0.15 – 0.43 | 0.517 | 0.15 | - 0.01 – 0.44 | 0.062 | 0.15 | 0.10 – 0.25 |
| Synchrony:FT | - 0.04 | - 0.30 – 0.28 | 0.783 | - 0.02 | - 0.19 – 0.16 | 0.788 | 0.04 | - 0.44 – 0.42 | 0.852 | 0.22 | - 0.12 – 0.52 | 0.184 | 0.05 | - 0.17 – 0.30 |
| Synchrony:SC | - 0.22 | - 0.47 – 0.01 | 0.057 | - 0.30 | -0.61 – - 0.04 | **0.019** | - 0.35 | - 0.85 – 0.12 | 0.145 | - 0.46 | -0.78 – - 0.20 | **0.001** | - 0.24 | -0.48 – - 0.03 |
| $R^2$ / $R^2$ adjusted | 0.308 / 0.241 | | | 0.250 / 0.175 | | | 0.122 / 0.042 | | | 0.300 / 0.238 | | | 0.221 / 0.148 | |

Note:

1. Confidence intervals (CI) and p-values are bootstrapped with 5,000 samples.
2. KC was used as a reference group.
3. Bs reflect the unstandardized regression coefficient

**Table S9.**

Bootstrapped regression for the relationship between mother and father behavioral synchrony moderated by birth group

| | Father-infant synchrony | | |
|---|---|---|---|
| *Predictors* | *B* | *CI* | *p* |
| Group FT | -0.38 | -1.97 – 1.20 | 0.632 |
| Group SC | 1.36 | 0.25 – 2.38 | **0.019** |
| Mother-infant synchrony | 0.29 | -0.06 – 0.57 | 0.088 |
| Mother-infant synchrony:group FT | 0.18 | -0.38 – 0.73 | 0.512 |
| Mother-infant synchrony:group SC | -0.60 | -1.05 – -0.09 | **0.022** |
| $R^2$ / $R^2$ adjusted | 0.210 / 0.140 | | |

Note:

1. Bootstrapped with 5,000 samples.

Simple slope analysis for the association between mother-infant and father-infant synchrony within each group revealed a significant positive correlation for FT ($p$<0.05), marginally significant for KC ($p$=0.09), but a negative correlation for SC ($p$=0.07).

**Table S10.**

Bootstrapped regression for the relationship between highest mother or father behavioral synchrony and dissimilarity levels moderated by birth group

| | **Joy-Sadness** | | | **Joy-Distress** | | | **Sadness-Distress** | | |
|---|---|---|---|---|---|---|---|---|---|
| *Predictors* | *B* | *CI* | *p* | *B* | *CI* | *p* | *B* | *CI* | *p* |
| Highest synchrony infancy | 0.18 | 0.13 – 0.28 | **0.001** | 0.20 | 0.10 – 0.29 | **0.003** | 0.18 | 0.06 – 0.34 | **0.006** |
| group FT [KC] | -0.10 | -0.51 – 0.41 | 0.657 | 0.24 | -0.24 – 0.81 | 0.363 | 0.48 | 0.08 – 0.90 | **0.019** |
| group SC [KC] | 0.52 | 0.18 – 0.86 | **0.006** | 0.22 | -0.30 – 0.60 | 0.354 | 0.21 | -0.36 – 0.79 | 0.433 |
| Highest synchrony infancy : group FC | 0.03 | -0.15 – 0.17 | 0.718 | -0.07 | -0.27 – 0.10 | 0.415 | -0.18 | -0.35 – -0.03 | **0.021** |
| Highest synchrony infancy : group SC | -0.19 | -0.33 – -0.04 | **0.018** | -0.03 | -0.19 – 0.20 | 0.735 | -0.03 | -0.26 – 0.22 | 0.814 |
| $R^2$ / $R^2$ adjusted | 0.377 / 0.321 | | | 0.298 / 0.235 | | | 0.284 / 0.219 | | |

Note:

1. Confidence intervals (CI) and p-values are bootstrapped with 5,000 samples.
2. KC was used as a reference group.
3. Bs reflect the unstandardized regression coefficient

**Table S11.**

Bootstrapped for the slopes of the relationship between highest mother-father behavioral synchrony and dissimilarity levels moderated by birth group

| | *Group* | *Est.* | *S.E.* | *t-value* | *CI - 2.5%* | *CI - 97.5%* | *P-value* |
|---|---|---|---|---|---|---|---|
| Joy - Sadness | FT | 0.215 | 0.051 | 4.211 | 0.115 | 0.316 | **0.000** |
| Joy - Sadness | KC | 0.186 | 0.055 | 3.360 | 0.078 | 0.295 | **0.001** |
| Joy - Sadness | SC | -0.004 | 0.062 | -0.063 | -0.126 | 0.118 | 0.950 |
| Joy - Distress | FT | 0.121 | 0.064 | 1.876 | -0.005 | 0.247 | 0.061 |
| Joy - Distress | KC | 0.197 | 0.071 | 2.795 | 0.059 | 0.336 | **0.005** |
| Joy - Distress | SC | 0.163 | 0.078 | 2.085 | 0.010 | 0.316 | **0.037** |
| Sadness-Distress | FT | 0.005 | 0.058 | 0.087 | -0.109 | 0.119 | 0.931 |
| Sadness-Distress | KC | 0.181 | 0.076 | 2.368 | 0.031 | 0.330 | **0.018** |
| Sadness-Distress | SC | 0.154 | 0.070 | 2.190 | 0.016 | 0.292 | **0.029** |

Note: Confidence intervals (CI) and p-values are bootstrapped with 5,000 samples.

**Table S12.**

Demographics

| | **KC** | **SC** | **FT** | *P-value* |
|---|---|---|---|---|
| **N** | 35 | 43 | 53 | |
| **Age (years) ±STD** | 18.63 ±0.84 | 18.67±0.94 | 21.29±2.10 | **<0.001** |
| **Sex** | 51.43% (18) Male; 48.57 (17) Female | 55.81% (24) Male; 44.19 (19) Female | 45.28% (17) Male; 54.72% (29) Female | 0.584 |
| **Dominant hand** | 71.429% (32) R; 22.86% (11) L | 74.419% (32) R; 25.59% (11) L | 88.67% (47) R; 9.43% (5) L | 0.102 |
| **Household Income above average (%)** | 41.18% (14) | 54.06% (20) | 82.69% (38) | **<0.001** |
| **Maternal level of education** | 91.18% (31) Academic | 74.36% (29) Academic | 89.13% (41) Academic | 0.080 |
| **Paternal level of education** | 64.71% (22) Academic | 69.70% (23) Academic | 86.96% (40) Academic | 0.051 |

**Table S13.**

Subjects excluded from analysis

| | KC | SC | FT | Total |
|---|---|---|---|---|
| **Scanned** | 35 | 44 | 58 | 137 |
| **Excluded due to movements** | 3 | 4 | 6 | 13 |
| **Misplaced in scanner** | 0 | 2 | 6 | 8 |
| **Structural abnormality** | 2 | 1 | 0 | 3 |
| **Technical problem with scanner** | 0 | 1 | 1 | 2 |
| **Other** | 1 | 2 | 5 | 8 |
| **Subjects Analyzed (%)** | 29 (82) | 34 (77) | 40 (68) | 103 (75) |
